# Microgradients in porosity and canal diameter in femur bone

**DOI:** 10.64898/2025.12.17.695028

**Authors:** Xiao Zhao, Xiaojun Yu, Swera Naz, Agila Zhussupova, Dilhan M Kalyon, Cevat Erisken

**Author notes:** These authors contributed equally to the work (Dilhan M Kalyon, Cevat Erisken). Corresponding Author. **Corresponding Author Address:** Dilhan M. Kalyon, Stevens Institute of Technology, 1 Castle Point Terrace, Hoboken, NJ, USA 07030; Cevat Erisken, Nazarbayev University, 53 Kabanbay Batyr Avenue, Astana, Kazakhstan 01000.

## Abstract

Bone exhibits hierarchical structural gradients that optimize mechanical performance and regenerative potential. Longitudinal and radial variations in cortical porosity and canal architecture influence load distribution, vascularization, and remodeling. Understanding these gradients is essential for designing scaffolds and implants that mimic native bone structure and function. This study quantified longitudinal and radial microgradients in cortical porosity and canal diameter along the rabbit femur and explored their implications for bone regeneration and repair implant design.

Rabbit femora were divided into proximal, mid-shaft, and distal regions. High-resolution micro-computed tomography quantified cortical thickness, porosity, and canal diameter along radial and longitudinal axes in micron-scale resolutions. Compressive mechanical testing of cortical slices determined local moduli, which were correlated with microstructural parameters to establish structure–function relationships.

Cortical thickness peaked at the mid-shaft and decreased toward both ends. Porosity and canal diameter increased radially toward the medullary cavity and longitudinally toward the bone ends. Upto 500 micron cortical thickness from the outer surface toward modullary cavity, porosity and canal diameter ranged, respectively, from ~5% and 40 µm at the mid-shaft to ~40% and 110 µm at the ends. At 750 micron cortical thickness, porosity and canal diameter ranged, respectively, from ~5% and 50 µm at the mid-shaft to ~80% and 200 µm at the ends. As expected, compressive moduli declined linearly with increasing porosity and canal size. The mid-shaft, with the lowest porosity and smallest canals, exhibited the highest modulus of around 15□MPa, which decreased to 5□MPa toward the ends.

The rabbit femur displays distinct longitudinal and radial microgradients in porosity and canal architecture that govern local stiffness. These gradients define structural benchmarks for designing functionally graded tissue engineering scaffolds and bone implants that replicate native tissue structure and stiffness transitions to promote osteoconduction, osteoinduction, osteogenesis in bone regeneration and improve osseointegration of bone implants.

## 1 Introduction

Bone quality encompasses more than just bone mineral density (BMD), incorporating critical microstructural features that determine mechanical competence and fracture resistance. ^1,2^ These include cortical thickness, porosity, and trabecular architecture. ^3,4^ While BMD remains a clinical standard, it often fails to predict fracture risk accurately, highlighting the importance of assessing bone microstructure ^5–7^. Bone quality changes are associated with conditions like osteoporosis and aging, contributing to increased fracture risk ^7^. Advanced imaging techniques, such as high-resolution CT and MRI, can provide detailed information about bone microarchitecture, potentially improving fracture risk prediction and treatment efficacy assessment ^2,8^. Understanding and quantifying bone microstructure is vital for enhancing diagnostic accuracy, developing therapeutic interventions, and informing biomimetic strategies in implant and scaffold design. ^3,6^

Cortical bone exhibits a complex hierarchical structure from the macroscale to the nanoscale, which underlies its mechanical properties and ability to resist fracture ^9,10^. This hierarchical organization spans from the diaphyseal shaft to microscopic osteons and vascular canals, down to the collagen-mineral matrix ^1,11^. Cortical bone accounts for approximately 80% of skeletal mass and plays a critical role in load-bearing, with its mechanical performance governed by mineral content and microstructural organization ^3,12^. Microstructural parameters such as porosity, canal diameter, and cortical thickness serve as essential indicators of bone health and mechanical resilience ^13^. The spatial distribution and density of vascular canals act as stress concentrators, influencing bone’s fatigue life ^14^. Quantitative assessment of cortical microarchitecture provides valuable insight into bone quality and structural integrity, especially in regions subject to high mechanical loads.

Recent studies have demonstrated that bone microarchitecture significantly influences its mechanical properties. Increased cortical porosity and enlarged vascular canals are associated with reduced bone strength ^1,15^. The intracortical canal network acts as a stress concentrator, affecting fracture risk ^16^. Accordingly, bone mineral density alone is insufficient to predict fracture risk, as microarchitectural parameters improve the prediction of mechanical behavior ^17^. Heterogeneity in material properties and microstructure contributes to fracture resistance and energy dissipation in bone tissue ^18,19^, while age-related increases in cortical porosity and Haversian canal size negatively impact bone’s ability to withstand mechanical loads^20^. Although porosity remains the primary determinant of cortical bone elasticity, other microarchitectural features also contribute at varying degrees ^21^. These findings highlight the importance of mapping compressive modulus alongside microstructural gradients, a more accurate and functional understanding of structure-function relationship can be achieved, enhancing the predictive value of microarchitectural assessment.

Spatial gradients in cortical thickness, porosity, and canal diameter are not merely an anatomical curiosity; they determine how host bone interacts with load-bearing implants such as femoral stems, tibial trays, and acetabular cups. Uniform titanium stems frequently over-stiffen the mid-shaft and under-support the more compliant metaphysis, promoting stress shielding and aseptic loosening. Implants with graded porosity, such as distally increasing axial gradients or inwardly increasing radial gradients, can improve osseointegration and reduce bone resorption ^22^. Designing implants with stiffness similar to cortical bone can minimize stress shielding and promote more physiological load transfer ^23,24^. Likewise, tissue-engineered scaffolds mimicking the natural bone structure, with a radial gradient of increasing porosity towards the center, have shown enhanced osteogenesis and vascularization. Studies demonstrate that higher porosity and larger pore canal diameters (>300 μm) promote bone ingrowth and vascularization in vivo ^25,26^. Biomimetic gradient scaffolds with fine internal pillars (~400 μm) and coarse external pillars (~800 μm) promote osteogenic differentiation and new bone formation ^27^. Quantitative mapping of these gradients in the present study therefore provides design targets for next-generation joint-replacement devices and bone/osteochondral grafts.

The rabbit femur is widely used as a preclinical model for studying bone structure and mechanics due to its similarities to human long bones ^28,29^. While not identical in scale or remodeling rates, the rabbit femur exhibits comparable cortical organization and loading environments to humans ^30,31^. The femoral shaft is particularly relevant for investigating microstructural features and biomechanics, with applications in orthopedic implant research ^32,33^. Rabbit models offer practical advantages including manageable bone size and established experimental protocols ^34^. However, it is important to note that no animal model perfectly replicates human bone, and understanding interspecies differences is crucial for accurate extrapolation of results ^29,35^. Despite some limitations, the rabbit femur remains a valuable platform for evaluating bone microstructure and biomechanics with translational relevance to human orthopedic applications.

Recent advancements in imaging techniques have significantly enhanced our ability to characterize bone microarchitecture and mechanical properties. Micro-computed tomography (micro-CT) allows high-resolution, three-dimensional quantification of bone structural parameters ^36,37^. This imaging modality, combined with sophisticated image processing, enables detailed analysis of bone architecture ^38,39^. Micro-CT resolution can reach 1 μm for ex vivo samples using synchrotron radiation ^40,41^, while in vivo imaging is limited to about 82 μm voxel size due to extensive radiation exposure concerns ^39^. Complementing imaging data, mechanical testing on bone samples provides critical information about functional performance ^42^. These techniques offer volumetric insights unattainable with traditional histology and have become standard tools for pre-clinical assessment of bone architecture during disease progression and treatment ^36,43^. The rabbit femur is particularly suitable for this dual mode characterization; its size permits both detailed imaging and consistent mechanical testing across multiple spatial planes. In contrast, the large diameter and density of human femurs often pose challenges for achieving the resolution and slice uniformity required for high-fidelity 3D structural-mechanical correlation, justifying the use of animal models in this context.

Despite growing interest in bone quality, most existing studies have focused either on global measures of bone mineral density or limited two-dimensional assessments of microstructure, without capturing the full spatial complexity of cortical bone. High-resolution, co-localized datasets that integrate three-dimensional microstructural metrics with corresponding mechanical properties remain scarce, particularly in the femoral shaft—a critical region for load-bearing and orthopedic applications. Furthermore, the radial and longitudinal gradients of features such as porosity and canal diameter is frequently overlooked, despite their likely influence on local mechanical behavior.

Few studies have quantitatively linked these microstructural parameters to compressive modulus in a spatially resolved manner. This study addresses these gaps by performing complete 3D profiling of cortical microarchitecture in the rabbit femoral shaft and systematically correlating these metrics with compressive properties across multiple spatial regions. The findings aim to advance our understanding of structure-function relationship of bone and inform the design of biomimetic materials with spatially optimized mechanical performance.

The primary aim of this study is to conduct a comprehensive three-dimensional characterization of cortical bone microarchitecture and its mechanical properties in the rabbit femoral shaft. Specifically, the objectives are: (1) to quantify gradual changes in porosity, canal diameter, and cortical thickness along both radial and longitudinal axes; (2) to measure compressive modulus in corresponding anatomical regions using mechanical testing on bone slices; and (3) to evaluate the correlations between microstructural parameters and localized mechanical properties. By integrating high-resolution imaging with direct biomechanical assessment, this study seeks to enhance the understanding of bone’s structure-function relationship and provide foundational data for the development of spatially informed designs. It is expected that the comprehensive and detailed characterization of the properties and structure of native bone that are presented here will allow for more realistic functionally graded design and fabrication of synthetic implants and scaffolds for tissue engineering.

## 2 Materials and Methods

Specimen preparation, instruments, and experimental design are demonstrated in Figure 1.

**Figure 1.**
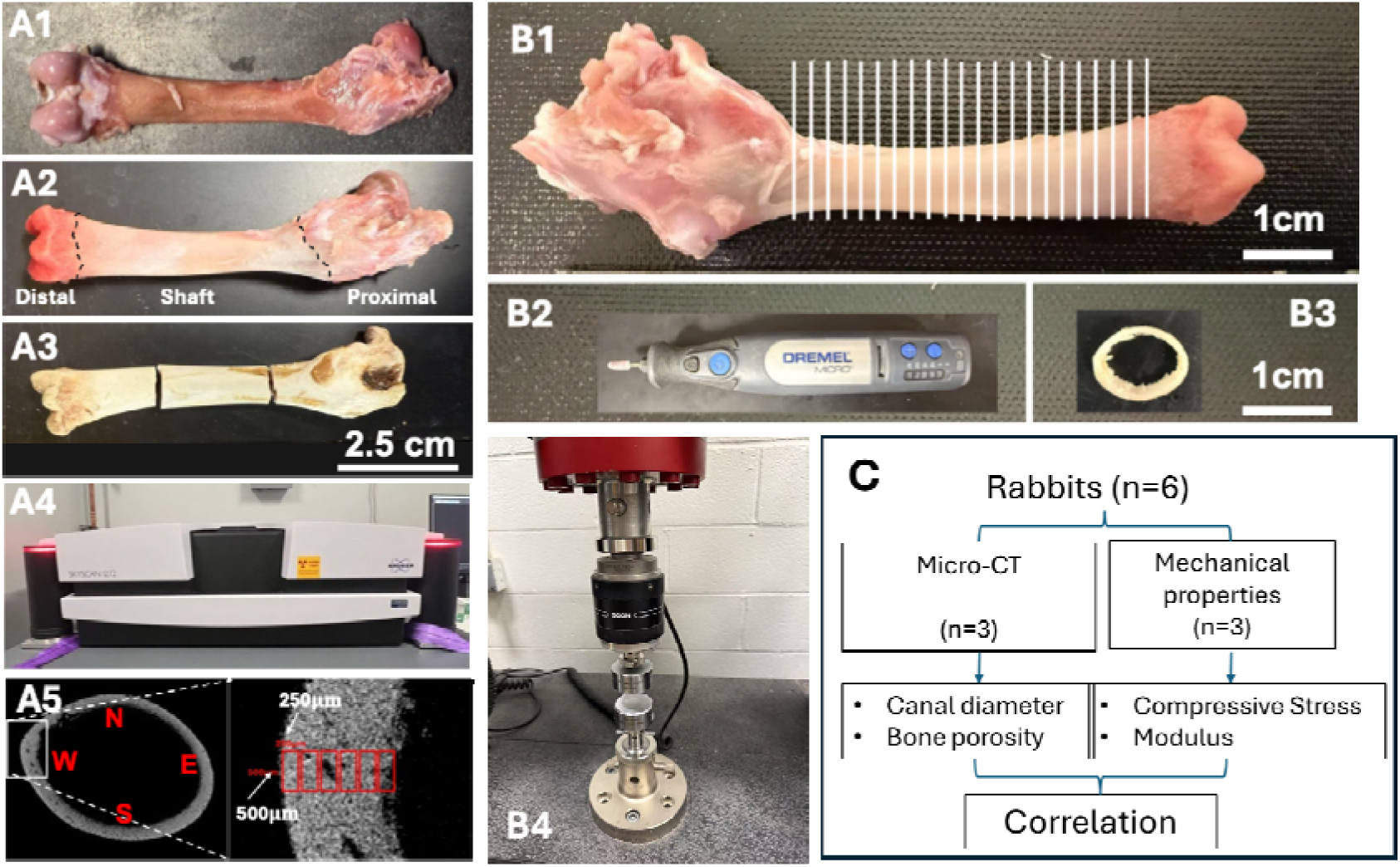
Specimen preparation, instruments, and experimental design. Femur bones (A1, A2) were harvested from rabbits, sectioned (A3) to fit into the sampling chamber of the micro-CT (A4), and analyzed in terms of pore and canal diameter by collecting data from different regions of the femur (A5). For mechanical characterization, the femurs were sliced in 2mm thicknesses (B1), processed (B2) to obtained uniform slice thickness (B3), and tested under compression (B4). A total of six rabbits were divided equally for micro-CT analysis and mechanical characterization (C).

### 2.1 Sample Preparation

Femoral bones were obtained from commercially sourced rabbits (Oryctolagus cuniculus) purchased from a local butcher shop (Jersey City, NJ). These rabbits (male and male) are meat-producing breeds (New Zealand White), slaughtered at an age of approximately 8–12 weeks.

Upon procurement, soft tissues surrounding the femur were carefully removed using a scalpel. The bones were then rinsed with phosphate-buffered saline (PBS) to remove residual debris and immediately stored at −20□°C to preserve structural integrity prior to analysis. All samples were thawed at room temperature for 2 hours before any subsequent imaging or mechanical testing. No chemical fixation or decalcification was applied, in order to preserve the native microarchitecture and mechanical properties of the bone tissue.

For imaging (n=3), following thawing at room temperature, each femur was segmented into three anatomical regions: the distal section containing the femoral condyles, the proximal section containing the trochanters, and the midshaft. The segmentation into distal, midshaft, and proximal regions was performed to analyze region-specific structural variations relevant to different loading conditions and enabled localized structural analysis across the major functional zones of the femur. After dissection, each bone section was thoroughly dried in a vacuum oven at 60□°C for 12 hours to ensure complete removal of residual moisture. This dehydration step was essential to prevent any water-related distortion or imaging artifacts and to maintain the structural integrity of the samples during subsequent handling. Care was taken to preserve the native morphology of each segment throughout the drying process.

For the compressive testing, fresh femoral bones were used (n=3). In this case, only the midshaft portion of the femur was utilized; the distal (condyle) and proximal (trochanter) ends were removed and excluded from analysis. The remaining shaft was evenly sectioned longitudinally into 2.5□mm-thick slices using a low-speed diamond saw (Dremel Multipro Model 395, Racine, WI). The longitudinal position of each slice along the femur was carefully recorded to preserve anatomical context. Each slice was then sanded down to a final thickness of 2.0□±□0.05□mm using a handheld drill fitted with a sanding attachment (Dremel Micro Model 8050, Mt. Prospect, IL). To avoid thermal damage or structural distortion, sanding was performed under continuous water cooling. This ensured the preservation of native mechanical properties and microarchitecture for downstream analysis.

### 2.2 Micro-CT Imaging and Analysis

Micro-computed tomography (micro-CT) imaging was conducted using the SkyScan 1272 system (Bruker microCT, Kontich, Belgium) equipped with a Photonic Science PS52 detector. Imaging was performed at a tube voltage of 100□kV and a current of 94□µA. A dual-filter configuration consisting of a 0.5□mm aluminum filter and a 0.038□mm copper filter was used to reduce beam hardening effects. The exposure time was set to 4000□ms per projection, with frame averaging set to three frames per projection. Scanning was carried out over a full 360° rotation with a rotation step of 0.4°, yielding a total of 900 projections per scan. The effective voxel size of the reconstructed images was 5.8□µm. Image reconstruction was performed using NRecon software (version 2.2.0.6, Bruker) with GPU acceleration enabled. Ring artifact correction was applied at level 12. Beam hardening correction and image smoothing were not applied in order to preserve the original grayscale distribution for subsequent quantitative analysis.

For the microstructural analysis, the 3D reconstructed model was sliced at 1□mm intervals. On each slice, the region of interest (ROI) was defined as a rectangular window measuring 500□×□250□µm^2^, positioned from the outer cortical surface toward the inner cortex. Measurements were taken at four orthogonal orientations—anterior, posterior, medial, and lateral—within each transverse micro-CT slice. These directional measurements were averaged to represent the radial structural profile at each longitudinal position, with data points collected at 250□µm, 500□µm, 750□µm, 1000□µm, etc., from the outer surface, continuing inward until no bone material remained within the ROI. Quantitative structural parameters extracted from each ROI included porosity (%), canal diameter (µm), and bone thickness (mm). All measurements were performed on the entire bone area within the defined ROI, restricted to the shaft region and excluding the distal and proximal femur, using calibrated image analysis protocols.

For the distal femur, proximal femur, and shaft segments, canal diameters were binned into defined size ranges, and the corresponding volume occupied by canals in each range was computed. The canal diameter distribution was expressed as the percentage volume of canals within a given diameter range relative to the total canal volume in the analyzed bone segment. This analysis allowed for comparative evaluation of canal architecture across distinct anatomical regions of the femur. Structural parameters derived from micro-CT imaging guided the interpretation of mechanical responses measured during subsequent compression tests.

### 2.3 Compression Testing

Compression testing was performed on bone slices using a uniaxial testing system (Instron 5980, Norwood, MA). Each specimen had a thickness of 2.0□±□0.05□mm and was derived from the femoral shaft region, excluding the distal and proximal ends. Prior to testing, all specimens were visually inspected to ensure parallel surfaces and absence of visible defects. Tests were conducted at room temperature with a crosshead displacement rate of 1□mm/min. No preload was applied. Load and displacement data were recorded continuously during each test and used to generate stress–strain curves.

Three mechanical parameters were extracted from the data: modulus 1, defined as the slope of the stress–strain curve in the first linear region; modulus 2, defined as the slope of the stress–strain curve in the second linear region; and maximum compressive strength, defined as the peak stress before failure. A total of three rabbits were used for this analysis, with multiple slices tested per animal to capture gradients along the femoral shaft. The exact number of slices per sample was recorded based on their anatomical position.

## 3 Results

### 3.1 Cortical Thickness Variation Along the Femur

To understand load-bearing adaptations, we first assessed variations in cortical bone thickness along the femoral shaft. Bone thickness was measured at 1□mm intervals along the longitudinal axis of the femur, starting from the distal end. The results demonstrated a non-uniform distribution of bone thickness along the length of the bone. As shown in Figure 2A, cortical thickness first decreased and then increased from the distal femur toward the midshaft, reaching a maximum in the central portion of the shaft, followed by a gradual decrease toward the proximal end.

**Figure 2.**
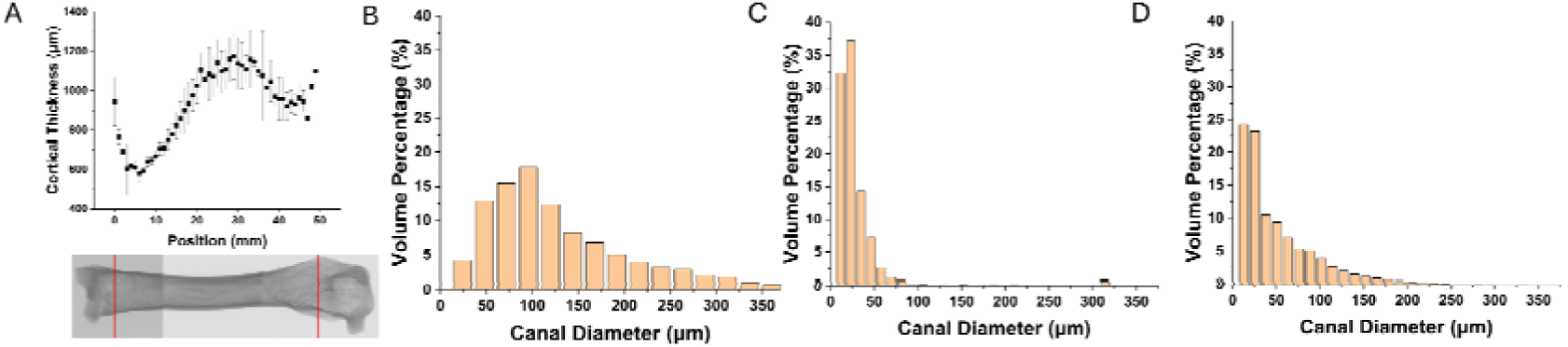
Change in cortical thickness with respect to distance from distal femur (A), and histogram of volume percent of the canal diameter for the distal femur (B), shaft (C), and proximal femur (D).

### 3.2 Regional Canal Diameter Distribution

Canal diameter distributions were quantified for the distal femur, shaft, and proximal femur to evaluate how intracortical vascular space varied regionally. Figures 2B–D show the volume percentage of canals distributed across diameter bins ranging from 0 to 375□µm. In the distal femur (Figure 2B), canal volume was primarily concentrated in the 50–200□µm range. The shaft region (Figure 2C) exhibited a narrower distribution, with the majority of canal volume confined to the 0–50□µm range. In the proximal femur (Figure 2D), canal diameters were mostly concentrated within the 0–100□µm range, with minimal volume beyond this threshold and a maximum diameter of approximately 250□µm.

The observed variability in canal diameter distributions across anatomical regions supports the selection of a fixed region of interest measuring 500□×□250□µm^2^ for micro-CT analysis. As demonstrated in Figure 2, canal diameters in the femoral shaft are predominantly below 50□µm, while the proximal and distal femur contain a broader distribution with canal diameters extending beyond 100□µm, and in some cases exceeding 250□µm, particularly in the distal region.

The chosen ROI dimensions allow for the consistent capture of the majority of canal structures present across all regions, while maintaining sufficient resolution for small-diameter canals in the shaft. Although some of the largest canals may partially fall outside the defined ROI, especially in the distal femur, the selected size offers a practical balance between resolution and anatomical coverage for comparative analysis. The distinct regional shifts in canal diameter suggested that void morphology varies not only in magnitude but also in spatial distribution; therefore, the following subsection maps porosity and mean canal diameter along both the femur’s length and its cortical thickness to integrate size metrics with overall void fraction.

### 3.3 Radial Gradients in Porosity and Canal Diameter

Porosity and canal diameter were analyzed along the radial direction (distance from the outer surface) of the femur using reconstructed micro-CT slices at 1 mm intervals in the longitudinal direction. Figure 3A1 presents a representative image of the full femur, with white dotted lines indicating the portion of the bone of which the porosity and canal diameter were measured as a function of the distance from the outer surface. Similar regional indicators are shown for the distal, shaft, and proximal segments in Figures 3B1, 3C1, and 3D1, respectively.

**Figure 3.**
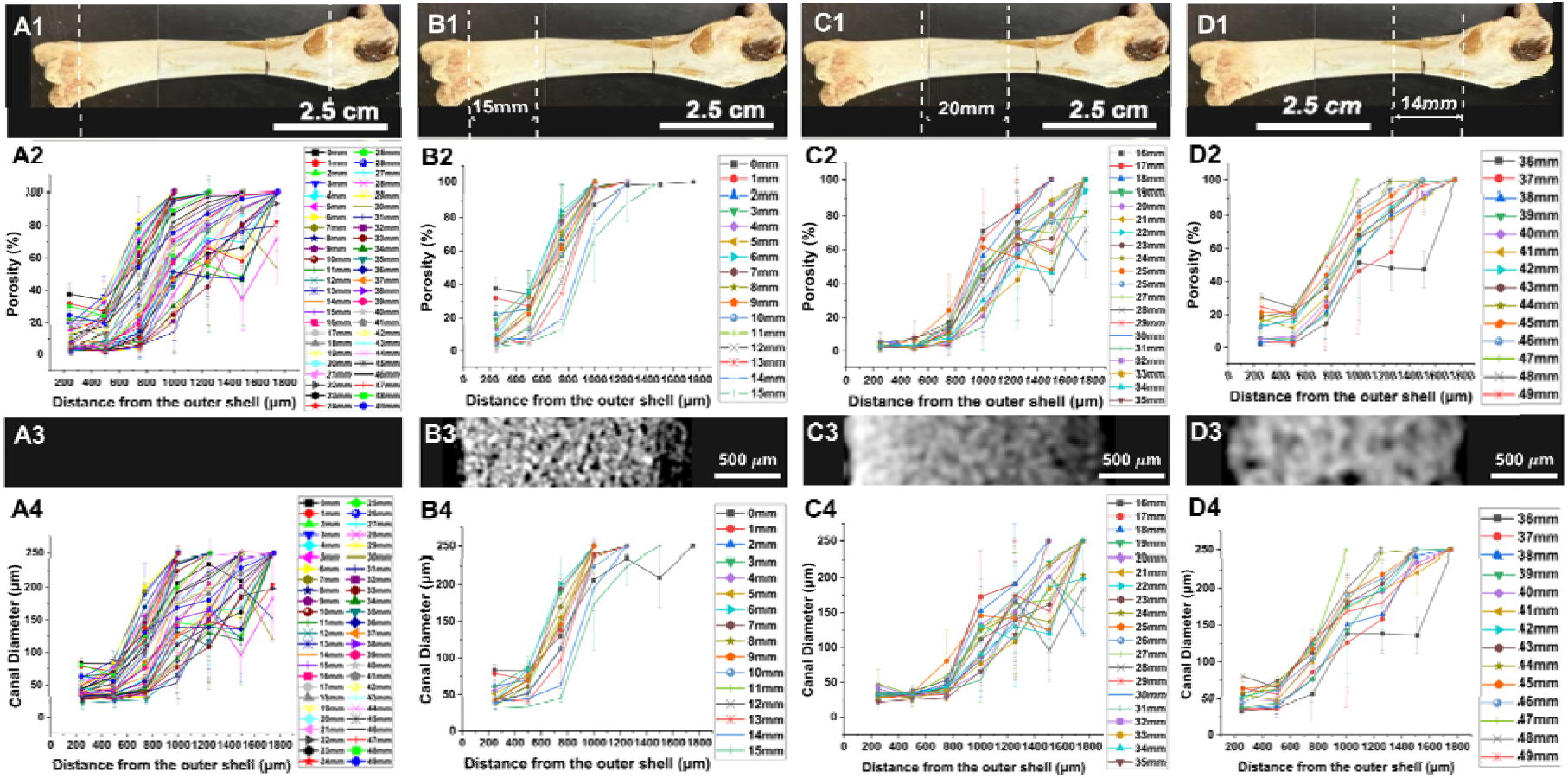
Change in porosity and canal diameter with respect to distance from the outer surface for the entire femur (A), distal end (B), shaft (C), and proximal end (D). The femur was scanned at 1mm intervals in the longitudinal direction for the segmented bones (B1, C1, D1) and merged to report the variation over the entire bone (A1). Porosity and canal diameter variation were measured from the scanned slices for the segmented bones (B2, C2, D2 and B4, C4, D4, respectively), and overlaid to obtain porosity change and canal diameter change over the entire scanned region (A2 and A4, respectively). Representative micro-CT images were provided from each segmented region (B3, C3, D3).

Regional measurements are detailed in Figures 3B2–D2 for porosity and Figures 3B4–D4 for canal diameter. In the distal femur, shaft, and proximal region, the porosity gradually increased from the outer surface towards the core of the bone. Representative transverse micro-CT images from each region are shown in Figures 3B3, 3C3, and 3D3. The distal (Figure 3B3) and proximal (Figure 3D3) regions show more porous bone with visibly larger and more numerous canals, while the shaft region (Figure 3C3) displays a denser structure with fewer and smaller canals, consistent with the quantitative porosity and canal diameter data.

**Figure 4.**
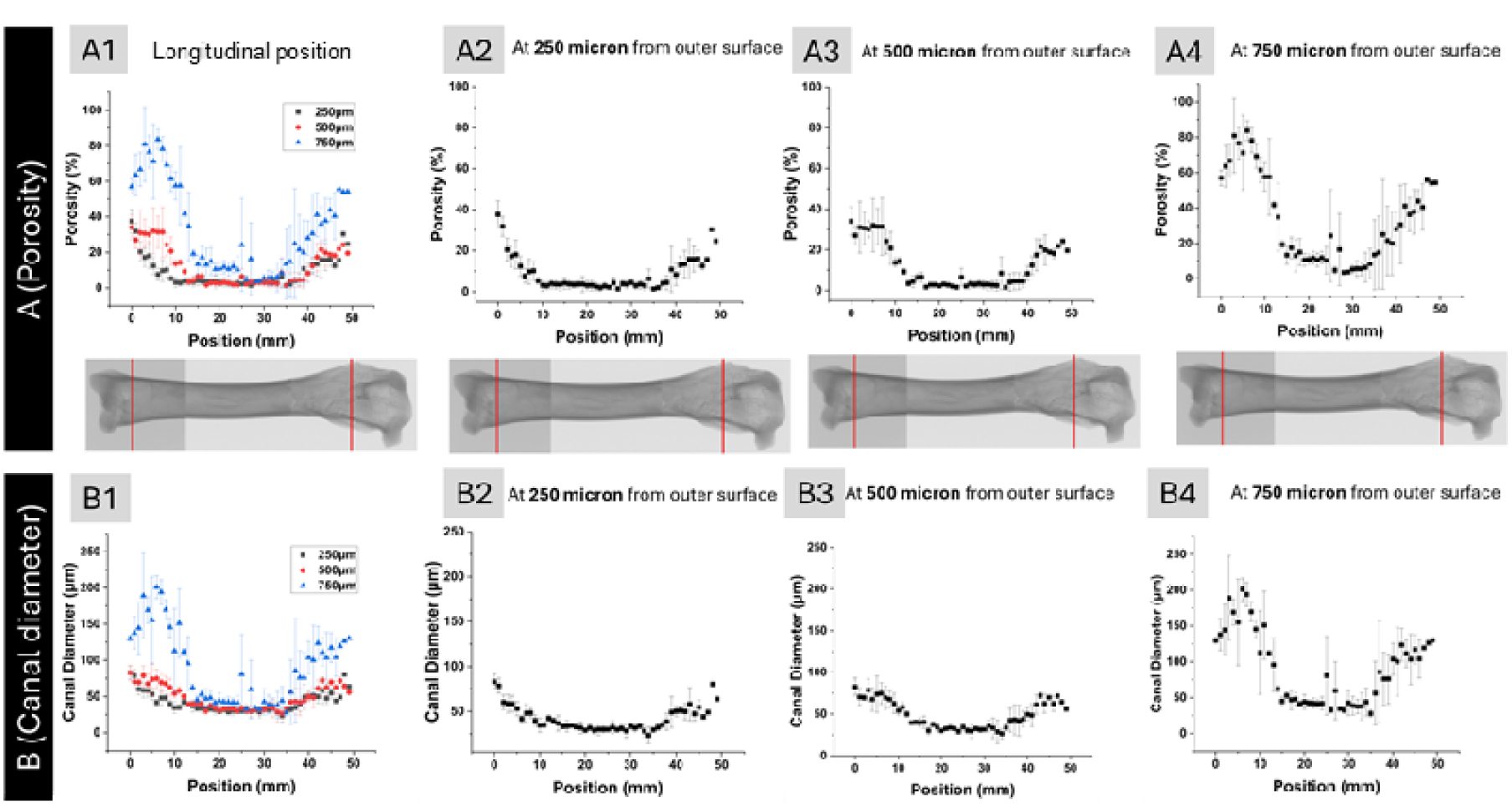
Porosity (A) and canal diameter (B) distribution in the femur as a function of longitudinal distance. Overlaid porosity distribution at 250, 500, and 750 μm distance from the outer surface (A1), and porosity distributions at 250 μm (A2), 500 μm (A3), and 750 μm (A4). Overlaid canal diameter distribution at 250, 500, and 750μm distance from the outer surface (B1), and canal diameter distributions at 250μm (B2), 500μm (B3), and 750μm (B4).

### 3.4 Longitudinal Microstructural Gradients

The data presented in Figure 4 were extracted from the same dataset used in Figure 3. For this analysis, porosity and canal diameter values were obtained specifically at 250, 500, and 750 µm from the outer surface and plotted against the longitudinal position along the femur.

The longitudinal overall trend in porosity is shown in Figure 4A for the entire bone length. Porosity values were higher in the distal and proximal regions and lower in the shaft. A similar trend was observed for canal diameter (Figure 3B), with reduced values in the shaft and elevated diameters toward both ends of the femur.

Figure 4A1 presents the overlay of porosity distributions at these three distances. The prof les shift with increasing distance, showing lower porosity at 250 µm and elevated porosity at greater distances from the outer surface. Individual porosity distributions at 250 µm, 500 µm, and 750 µm are shown in Figures 4A2, A3, and A4, respectively. Porosity at 250 µm was relatively low and exhibited a narrow distribution, while porosity at 500 µm and 750 µm increased in both magnitude and spread, indicating a gradient in structural void content across the cortical thickness.

Canal diameter distributions across the same distances from the outer surface are shown in Figure 4B1 as an overlay, with individual profiles at 250□µm, 500□µm, and 750□µm presented in Figures 4B2, B3, and B4, respectively. At 250□µm, canal diameter along the femur was concentrated within the 0–75□µm range, with limited variation between anatomical regions. At 500□µm, the overall range remained similar, with most values still falling below 75□µm; however, slight local increases were observed in the distal and proximal regions. In contrast, at 750□µm, canal diameter increased noticeably in the distal and proximal regions, reaching values up to 200□µm, while the shaft remained confined to a lower range comparable to that at 250□µm and 500□µm. These data indicate that substantial enlargement of canals with respect to longitudinal position primarily occurs at greater distances from the outer surface.

Figure 5 presents the correlation between porosity and canal diameter at 250, 500, and 750□µm from the outer surface. This plot was generated using the same dataset shown in Figure 4 by pairing porosity and canal diameter values at corresponding positions along the femur. A linear relationship was observed at each distance, with comparable slopes across the three curves. While the overall trend was consistent, the curve at 750□µm extended to higher values than those at 250 and 500□µm. Specifically, porosity reached approximately 80% and was associated with canal diameters up to 200□µm in the distal and proximal regions. This extended range at 750□µm reflects the localized increase in both canal size and void volume deeper within the bone at those positions.

**Figure 5.**
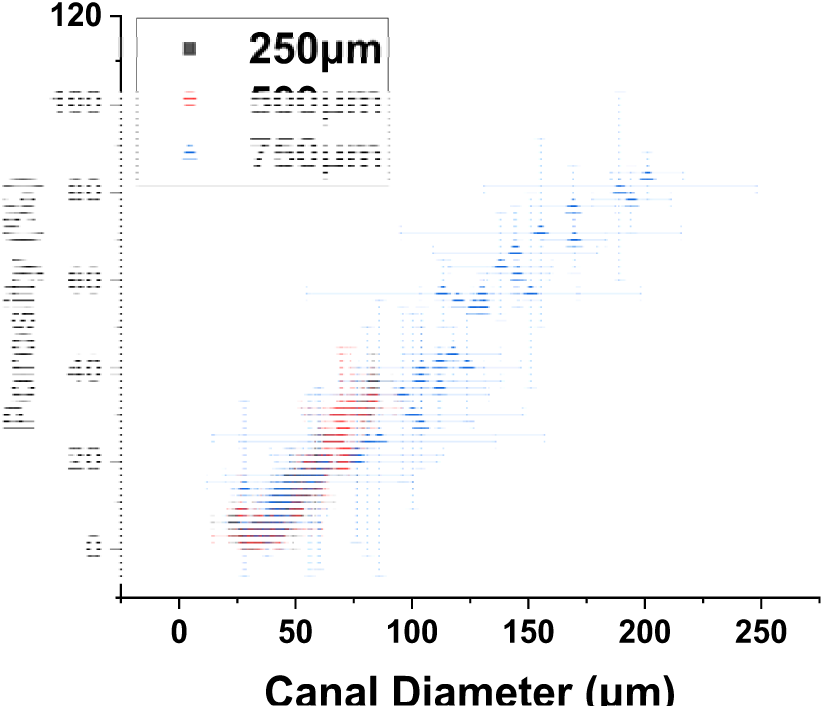
Correlation of porosity and canal diameter at 250, 500, and 750 μm distance from the outer surface.

Quantifying how porosity and canal diameter evolve from the outer to the inner cortex raised the question of whether these microstructural changes translate into local mechanical behavior; accordingly, the next section relates the radial data to position-matched compressive moduli.

### 3.5 Mechanical Properties and Microstructure Correlation

Relationships between microstructural parameters and mechanical properties were evaluated to confirm the role of microarchitecture in stiffness. Correlations between bone microstructure and compressive properties were assessed by relating porosity and canal diameter to modulus 1 and modulus 2. Figure 6A1 shows the relationship between porosity and modulus 1. As porosity increased from approximately 5% to 40%, modulus 1 decreased from about 15□MPa to 5□MPa. A similar inverse correlation was observed between porosity and modulus 2 in Figure 6A2. Over the same porosity range (5% to 40%), modulus 2 declined from approximately 1.4□MPa to 0.7□MPa.

Figures 6B1 and 6B2 show the correlations between average canal diameter and modulus 1 and modulus 2, respectively. As canal diameter increased from approximately 20□µm to 110□µm, both modulus 1 and modulus 2 exhibited decreasing trends. Modulus 1 declined from around 15□MPa to approximately 5□MPa over this range. Similarly, modulus 2 decreased from about 1.4□MPa to 0.7□MPa. Although the data were more scattered compared to the porosity-based correlations, the overall negative association between canal diameter and compressive stiffness was consistently observed in both metrics.

Figure 6C presents compressive stress–strain curves for all femur slices subjected to mechanical testing. The legends indicate the slice numbers, which correspond to their longitudinal positions along the femur. Each curve exhibited two distinct linear regions. The slope of the first linear region was used to calculate modulus 1, while the slope of the second linear region was used to calculate modulus 2, as described in the methods section. The presence of two linear regions across all tested slices demonstrates the consistency of this mechanical behavior throughout the femur. While the point correlations confirm that stiffness diminishes with increasing void volume and canal size, a continuous profile is required to visualise how structural and mechanical properties co-vary along the entire femoral shaft; the subsequent analysis provides this integrated longitudinal map.

**Figure 6.**
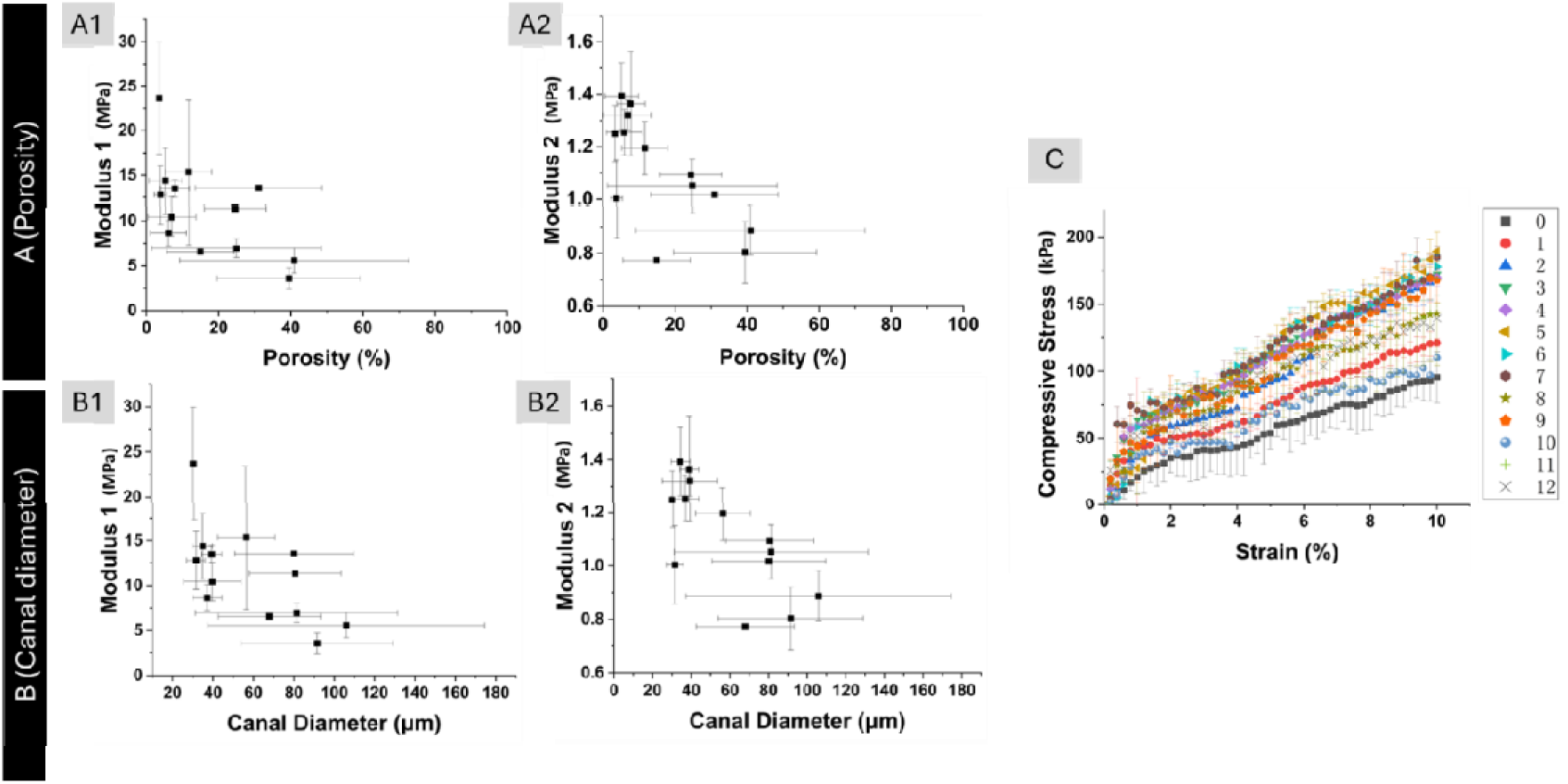
Correlation of modulus 1 and porosity (A1), modulus 2 and porosity (A2). Correlation of modulus 1 and canal diameter (B2), modulus 2 and canal diameter (B2). Compressive stress and strain curve for mechanical compression test (C). In (C), the legend shows the number of sections with ~2mm thickness.

### 3.6 Longitudinal Variation of Mechanical and Structural Properties

Finally, structural and mechanical profiles were integrated to reveal spatial coordination along the femur shaft. Figure 7 presents the variation of microstructural and mechanical parameters along the longitudinal axis of the femur. Average porosity and average canal diameter trends are shown in Figures 7A and 7B, respectively. Average porosity values were lowest in the shaft region, ranging between 2% and 5%, and increased progressively toward the distal and proximal ends, reaching up to approximately 40%. Average canal diameter exhibited a similar longitudinal pattern, with values between 30 and 40□µm in the shaft and increasing to as high as 110□µm at the bone ends. These trends are consistent with the regional and radial analyses presented in earlier sections.

**Figure 7.**
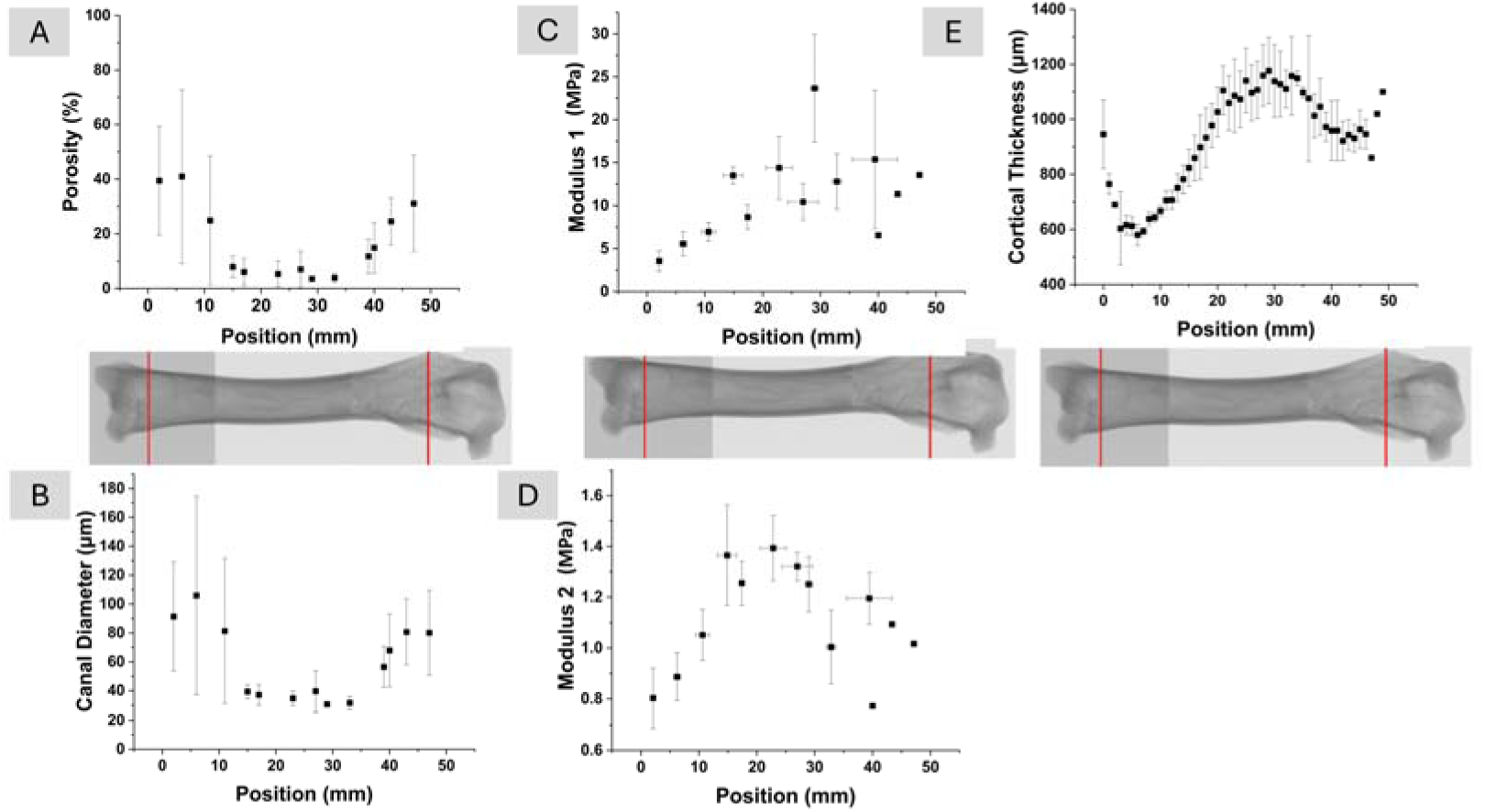
Porosity (A) and canal diameter (B) distribution in the femur as a function of the position. Correlation of modulus 1 and position (C), modulus 2 and position (D). Longitudinal variation in bone thickness (E).

Figures 7C and 7D show the variation of modulus 1 and modulus 2 along the longitudinal axis of the femur. Both modulus values peaked in the shaft region and decreased toward the bone ends. Modulus 1 ranged from approximately 5□kPa at the distal end (0□mm), increased to around 25□kPa in the central shaft, and then declined to about 10□kPa at the proximal end (50□mm). Modulus 2 followed a similar trend, rising from approximately 0.7□kPa at the distal end to 1.4□kPa in the shaft and decreasing to around 1.0□kPa at the proximal end. These longitudinal profiles reflect a central zone of greater compressive stiffness, consistent with the lower porosity and smaller canal diameter observed in the shaft region.

Figure 7E shows the longitudinal variation in bone thickness. The overall pattern of bone thickness closely follows the trends observed for modulus 1 and modulus 2 in Figures 7C and 7D. Bone thickness increased from the distal end toward the shaft and then gradually decreased toward the proximal end. This progression corresponds with the region of highest compressive stiffness and lowest porosity along the shaft, as shown in Figures 7A and 7B. The spatial alignment of bone thickness with structural and mechanical parameters suggests a consistent anatomical transition along the femur.

## 4 Discussion

Our micro-CT analysis revealed that cortical thickness was highest at the femoral mid-shaft and tapered toward both proximal and distal ends. This region is known to carry the largest bending and axial loads during routine gait, so the extra thickness likely reinforces the bone against fracture ^44–46^. The thinner cortices near the bone ends may represent an anatomical compromise that provides space for blood vessels and allows a gradual transition to trabecular bone at the metaphysis ^47,48^. Previously, Pazzaglia et al. studied New Zeeland rabbits and found no significant difference between the cortical thicknesses of distal and proximal femur.^49^ Overall, these results support that region-specific variations in cortical thickness are a key structural factor governing the mechanical competence of long bones.

Following the thickness analysis, micro-CT imaging showed that vascular canals were widest in the distal cortex (50–200 µm), smallest at the mid-shaft (< 50 µm), and intermediate proximally. This regional pattern means that screws or rods of implants pressed into the distal cortex engage a more porous, less stiff substrate than at the shaft, creating a stiffness mismatch that has been linked to early screw loosening and plate-end failure in clinical studies ^22,50,51^. A mismatch between the implants and the native bone tissue could additionally place unbalanced loads over the knee joint, further damaging the joint components, including the anterior cruciate ligament (ACL). This unbalanced loading may tear/rupture the ACL, leading to joint instability ^52–54^. Modern plates and scaffolds are usually uniform in density; reproducing the larger distal canals while preserving the tighter mid-shaft architecture could reduce such failures and speed blood-vessel ingrowth during regeneration. Our quantitative canal-diameter map therefore offers practical dimensions for graded implants and additively manufactured scaffolds that aim to match native cortical structure along the bone length.

Both longitudinally and radially, porosity and canal diameter increased notably toward the femoral ends. Porosity and mean canal diameter were roughly three-times higher at the femoral ends (~ 28 % porosity; ~ 110 µm canals) than at the mid-shaft (~ 9 %; ~ 40 µm). These longitudinal gradients lower stiffness where loads are smaller and boost blood supply and remodeling capacity near joints, but it also means that uniform metal plates or nails over-stiffen the porous ends and can concentrate stresses that lead to screw loosening or end-fracture. Additively manufactured implants and scaffolds that grade pore size—larger at the ends, denser at mid-shaft—would better share load with the host bone ^55^ and speed vascular invasion during healing ^56–58^.

Moving from lengthwise patterns to the radial direction, porosity and canal diameter rose steadily from the periosteal surface toward the medullary cavity, reaching values of about 80 % porosity and ~200 µm canals in the inner cortex of distal and proximal regions. The radial gradient in porosity and canal diameter, increasing notably from the periosteal surface inward, indicates structural adaptation that might prevent full-thickness fracture propagation ^59^. Layered implants or scaffold struts that are denser externally and more open internally would better match native load transfer and speed marrow-side vascular invasion ^60^; present devices rarely achieve such through-thickness grading.

Building on the spatial gradients already described, our findings showed that both modulus 1 (initial modulus) and modulus 2 (secondary modulus) dropped wherever porosity or mean canal diameter increased: denser regions with smaller canals were stiffer, while more porous regions with larger canals were more compliant. To our knowledge, this is the first study to chart modulus 1 and modulus 2 against porosity and canal diameter across the full shaft of the rabbit femur model, giving a practical way to estimate position-specific stiffness with regard to microstructural properties. Together, these findings support microarchitecture, rather than bulk mineral density, governs cortical mechanics and supply reference values for future comparative or modelling work ^1,15,16,61,62^.

Our integrated analysis clearly showed mid-shaft as the zone of highest structural robustness and stiffness. Modulus 1 and modulus 2 were highest at the femoral mid-shaft and declined by roughly 40 % toward both distal and proximal ends, matching the longitudinal shifts in thickness, porosity, and canal size described earlier. This alignment shows that the femur gradually softens where loads are lower, a strategy that avoids stress peaks and saves material. Conventional plates and intramedullary rods are uniform in stiffness; by over-supporting the naturally stiff mid-shaft and under-matching the more flexible ends, they can shift load to screw holes and promote plate-end failure ^22,63^. Functionally graded implants that are rigid at the mid-shaft and progressively more compliant toward the ends would better track the native modulus profile as would additively manufactured scaffolds printed with a similar stiffness gradient to guide regeneration ^64,65^.

The spatial maps of bone thickness, porosity, canal diameter, and modulus presented in this chapter offer quantitative targets for the next generation of total hip and knee replacement devices and for tissue-engineered osteochondral grafts ^66–68^. Metallic stems, cups, and tibial trays manufactured from uniform Ti-6Al-4V or Co–Cr alloys typically possess elastic moduli far exceeding those of host bone, particularly at the metaphyseal ends, thereby redistributing load and promoting stress shielding ^22^. By documenting a pronounced axial drop in cortical stiffness from the femoral mid-shaft toward both the distal and proximal regions concomitant with a marked rise in porosity and canal diameter, the present study defines the magnitude and length-scale of the gradient that graded stems or trays must reproduce to maintain physiological load transfer and mitigate aseptic loosening.

Axial grading alone, however, cannot resolve interfacial mismatch if radial heterogeneity is ignored. The observed progressive increase in porosity and canal diameter from the outer cortex toward the medullary cavity demonstrates that cortical bone functions as a dense shell surrounding a more compliant, highly vascularized core. Functionally graded scaffolds with seamless gradient that emulate this radial transition can therefore couple high interfacial stiffness and wear resistance at the surface with enhanced vascular and cellular infiltration internally ^27,69^. Such bimodal designs align with emerging porous-metal and polymer-ceramic composites that report accelerated osseointegration when inner canal diameters exceed one hundred micrometers ^25^.

Beyond empirical design, the strong correlations quantified here between local modulus and structural metrics such as porosity and canal diameter establish data-driven rules for assigning position-dependent material properties in the analysis of implants and grafts. Incorporating these correlations allows simulations to capture the gradual load hand-off from implant to host tissue, providing a foundation for optimizing regional implant density, structure thickness, and material composition prior to manufacturing. Earlier, a strong dependence of modulus on porosity was also determined at smaller length scales.^70^

Our approach combined high micro-CT with location-matched compression tests, allowing us to link local modulus values to canal diameter and porosity with millimeter-scale precision. Although the rabbit model provides valuable insights due to practical advantages, higher turnover rates and anatomical differences limit direct translation of absolute thresholds to human bones, highlighting the need for cautious extrapolation. Additional limitations include the modest sample size, removal of periosteum during preparation (which may alter hydration), and the use of uniaxial compression rather than bending, all of which could influence absolute modulus values. Even so, the thorough 3D study and profiling of rabbit femur microstructural and biomechanical properties strengthen confidence in the positional patterns reported here and provide a clear baseline for future graded-implant and scaffold studies.

Future studies should extend these findings to larger animal or human cadaveric models, evaluate changes with aging and disease, and use these refined data sets to directly guide the design of implants and scaffolds optimized for clinical translation. Second, longitudinal micro-CT and matched mechanical testing in ageing or osteoporotic rabbits could track how these spatial patterns evolve with disease or time and establish thresholds for intervention. Third, the quantitative maps of thickness, porosity, canal size, and modulus reported here can be integrated into the design loop for implants and scaffolds for tissue engineering, enabling graded biomaterials that better match native stiffness and vascular requirements than today’s uniform devices. Functionally graded materials (FGMs), i.e., composites characterized by spatially varying properties that transition continuously or discretely across dimensions ^15,16,71–74^ can be used.

Such materials can be designed to address the limitations inherent in homogeneous systems, particularly at interfaces where abrupt property differences can lead to mechanical failure or poor biological integration ^16,17^. FGMs enable gradual changes in composition, microstructure, or porosity, facilitating improved load transfer, stress distribution, and material compatibility in structural and biomedical applications ^15,18,19,74^.

The osteochondral interface, which connects articular cartilage to the underlying subchondral bone, is a prototypical example of a biological FGM. It transitions from a viscoelastic, avascular cartilage layer to a highly mineralized bone layer, reflecting a continuous increase in mechanical stiffness across the interface ^22,26^. Mimicking this gradient would be essential to engineering grafts that perform comparably to native tissue under physiological loading conditions ^23,25^.

An additional recent focus in biomaterials research has been on integrating fabrication techniques that are industrially relevant, such as extrusion-based bioprinting, hybrid extrusion and electrospinning, coextrusion for multiple layers, hybrid extrusion and spiral winding, melt electrowriting to develop FGMs with fine-tuned gradients in mineral content and mechanical properties ^66,75–87^. Such technologies have the potential to facilitate seamless transitions between distinct biomaterial phases and can be instrumental in the development of osteochondral scaffolds that closely mimic the native interface ^88–90^.

Lastly, while the rabbit model provides rapid healing and tractable size, its higher turnover rate and thinner cortex mean that absolute thresholds may differ from those in larger animals or in humans. Extending the present mapping to ovine or cadaveric femoral shafts, and tracking how the gradients shift with age, disease, and cyclic loading, will refine the design window for clinical devices. Integrating our micro-CT–derived modulus relationships into finite-element simulations and iterative implant prototyping will further accelerate translation.

## 5 Conclusion

This work set out to create a detailed, spatially resolved picture of structure–function relationships along the rabbit femoral shaft and to translate those insights into design criteria for next-generation orthopedic implants and tissue-engineered scaffolds. By pairing high-resolution micro-CT imaging with location-matched compression testing, we mapped bone thickness, porosity, vascular-canal diameter, and local modulus at millimeter-scale precision, producing, to the best of our knowledge, the first full-wall dataset of its kind for any long bone in the rabbit model.

Three consistent patterns emerged. First, bone thickness peaked at mid-shaft and tapered toward both ends, aligning with the region of greatest habitual bending and axial load. Second, porosity and mean canal diameter rose not only from shaft to ends but also from the outer surface toward the medullary cavity, creating combined longitudinal and radial gradients in void fraction. Third, both the initial and post-yield moduli fell wherever porosity or canal size increased, demonstrating that microarchitecture—rather than bulk mineral density—governs cortical stiffness. Together, these findings reveal a shaft that is simultaneously thick, dense, and stiff, flanked by ends that are thinner, more porous, and mechanically compliant. Translating these observations to practice, the numeric profiles reported here offer concrete targets for functionally graded plates, intramedullary rods, and additively manufactured scaffolds. Implants that preserve a stiff mid-shaft segment while gradually reducing stiffness and increasing permeability toward the bone ends—and toward the marrow side—could mitigate screw loosening, stress shielding, and delayed union. Likewise, scaffolds patterned with smaller, load-bearing pores at mid-shaft and larger, angiogenic pores distally and medially should better balance mechanical support with vascular access during regeneration.

## Data Availability

All the data available were submitted with this manuscript.

## Clinical trial number

Not applicable. The bone tissues were obtained from animals slaughtered for food production.

## Author contribution

XZ, SN, AZ conducted experiments; XZ drafted the manuscript; XY, DMK, CE edited the manuscript; XY, DMK provided materials; CE provided funding.

